# Individual differences in the effects of salience and reward on impulse control and action selection

**DOI:** 10.1101/2022.12.23.521803

**Authors:** I. Schutte, D.J.L.G. Schutter, J.L. Kenemans

## Abstract

Impulse control and adequate decision making are vital functions when it comes to detection and adherence to societal rules, especially in critical circumstances such as the Covid pandemic. In the current study we tested the hypothesis that increasing the salience of environmental cues would be most effective in improving impulse control, as assessed in a stop-signal task, in subjects with low environmental susceptibility as indexed by low pre-stimulus EEG alpha power. In addition, we anticipated that an external-reward intervention improves performance during a Go/No go task, especially in individuals with low task-induced motivation as indexed by low theta/ beta power ratios. High salience of stop signals enhanced stopping performance but there was no difference in responsivity to the salience intervention between participants with high and low EEG alpha power. Individuals with low theta/ beta power ratios responded more accurately when rewards were at stake. Together these results suggest that increasing the salience of external cues may help impulse control in general, whereas the effectiveness of external-reward interventions is higher in individuals with low task-induced motivation.

## 1. Introduction

Impulsivity can be defined as a range of actions that are “poorly conceived, prematurely expressed, unduly risky, or inappropriate to the situation, and that often result in undesirable outcomes” (Daruna & Barnes, 1993). The ability to control impulsive actions has important implications for social and economic decision-making in daily life, but pronounced individual differences exist.

The ability to keep control over impulses is, for example, important in situations where individuals have to follow rules in order to prevent harm to themselves or others, such as the physical distancing rules in the context of the Covid-19 pandemic. Humans have an innate tendency to engage in close social and physical contact and adherence to physical distancing rules may therefore be difficult. Some individuals know the rule(s) and perhaps intend to adhere, but nonetheless fail to comply on many occasions.

This raises the question of whether impulsive tendencies can be influenced. In the current study, we considered two possible interventions that may enhance impulse control: increasing the salience of external cues that signal the need for impulse control, and rewarding fast and accurate action selection.

Several studies including a previous study in our lab have shown that increasing the salience of external cues that signal the need for impulse control enhances stopping competence (Blizzard, Fierro-Rojas, & Fallah, 2017; Kenemans, Schutte, Van Bijnen, & Logemann, 2022; Montanari, Giamundo, Brunamonti, Ferraina, & Pani, 2017; Van Der Schoot, Licht, Horsley, & Sergeant, 2005), the generally accepted objective index of impulse control (Kenemans, 2015). Furthermore, it has been shown that rewards affect stopping competence (Boehler, Schevernels, Hopf, Stoppel, & Krebs, 2014) as well as task performance in general (Schutte, Heitland, & Kenemans, 2019). However, how individual characteristics can contribute to differences in responsivity to such interventions remains unclear.

It has been shown that enhanced alpha power in the electroencephalogram (EEG) is associated with reduced susceptibility to potentially critical external signals (Dockree et al., 2017; O’Connell et al., 2009). This may indicate that individuals with high alpha power are less able to attend to and/or process environmental cues, including cues that signal the need for impulse control.

The ratio between low-frequency EEG theta and high-frequency EEG beta power has been put forward as a trait-like index of reward sensitivity (Massar, Kenemans, & Schutter, 2014; Schutter & Kenemans, 2022; Schutter & Van Honk, 2005) and has been found to explain unique variance in reward-related task performance (Schutte, Kenemans, & Schutter, 2017). Increased theta/ beta power ratios were for example associated with enhanced reward-learning (Massar et al., 2014) and risky decision-making and reward-seeking during the Iowa gambling task (Massar et al., 2014; Schutter & Van Honk, 2005). A previous study revealed that individuals with higher theta/ beta power ratios during resting state were also better at stopping (Lansbergen, Schutter, & Kenemans, 2007). This observation was explained in terms of these individuals being driven by increased task-induced motivation to maximize stopping performance.

In the current study, we used existing data sets to relate EEG indices of environmental susceptibility and reward sensitivity to stopping competence, and reward-induced enhancement of action selection in a Go/Nogo context. The rationale behind the present work is that EEG alpha and theta/ beta power assessments may provide useful means to examine individual states of susceptibility and reward sensitivity. In turn, this enabled us to assess individual differences in responsivity to interventions regarding the salience of withhold-cues and regarding reward manipulation.

Specifically, we addressed five hypotheses: 1) High susceptibility/ low alpha predicts better impulse control. 2) Enhanced stop-signal salience induces better impulse control. 3) Enhanced salience induces better impulse control in specifically low-susceptibility/ high alpha individuals. 4) Introducing a reward induces more accurate and faster action selection. 5) Introducing a reward induces more accurate and faster action selection specifically in individuals with low levels of task-induced motivation (i.e., individuals with low theta/ beta power ratios). Furthermore, a secondary aim was to replicate our previous finding that theta/ beta power ratio during rest is associated with better stopping competence (hypothesis 6).

A combination of stopping performance data under conditions of low and high stop-signal salience and stop task-related EEG data allowed us to analyze whether low alpha power preceding the stop signal is correlated with better stopping competence as indexed by stop signal reaction times (SSRTs; hypothesis 1). In addition, it enables to examine the effect of salience on SSRT, and whether this effects differs for low-versus high-alpha-power groups (hypotheses 2 and 3, respectively). If anything, this relationship was expected to particularly manifest in a condition in which high trait susceptibility is presumably most beneficial. We, therefore, analyzed these data for a condition during which the stop signal is low-salient and subsequently examined whether salience enhancement results in better stopping performance, and mostly so in individuals with high alpha power preceding low-salient stop signals.

A combination of performance data in a cued-go/nogo reaction-time task (Schutte et al., 2019), under conditions of reward versus no reward, and resting-state EEG data allowed us to address hypothesis 4 and 5: Does reward induce better action selection, and mostly so in participants with low theta/ beta power ratios? Finally, a third data set consisting of stopping performance and resting-state EEG allowed us to attempt to replicate our previous finding (Lansbergen et al., 2007) of a positive relation between theta/ beta power ratio during rest and stopping competence (hypothesis 6).

## 1. Methods

### 2.1 Subjects and datasets

For hypotheses 1, 2, and 3 we used stop-signal EEG and stop-signal performance data from one previously collected sample (n = 30; (Kenemans et al., 2022)). For hypothesis 4 and 5 we combined previously collected data from two samples (n = 48 (Schutte et al., 2019); 1 subject lost, see 2.5.4, and n = 8 (Schutte, Deschamps, van Harten, & Kenemans, 2020)). For hypothesis 6 (replication of the correlation between theta/ beta power ratio and stopping competence) auditory stop-signal data and resting state EEG data from one previously collected sample (n = 52; (Massar, 2012); 1 subject lost, see 2.5.4) were used. Table 1 presents an overview of the research questions and associated subject samples, and the descriptive statistics.

**Table 1.** Descriptive **statistics**

| Hypothesis | Used data set(s) | N analyzed | Gender (f/m) | Age ($\bar{y} \pm \text{sd}$) |
| --- | --- | --- | --- | --- |
| anticipated relationship pre-stim alpha, stopping competence, and salience (hypotheses 1, 2, 3) | Kenemans et al., 2022 | 30 | 24/6 | 23 ($\pm 3.3$) |
| anticipated relationship between resting state theta/beta power ratio and reward-related performance (hypothesis 4, 5) | Schutte et al., 2019;<br>Schutte et al., 2020 | 55 | 36/19 | 23 ($\pm 3.9$) |
| anticipated relationship between resting state theta/beta power ratio and stopping competence (replication; hypothesis 6) | Massar, 2012 | 51 | 27/24 | 22 ($\pm 3.1$) |

All subjects were healthy and reported normal or corrected-to-normal vision. Subjects who participated in the stop task experiments additionally reported normal hearing. All were recruited mainly among the student population. Possible other in and exclusion criteria are listed in the manuscripts mentioned above. Subjects were asked for permission to re-use their data. The study was approved by the local ethics committee of the faculty of Social and Behavioral Sciences of Utrecht University.

### 2.2 Procedure

#### 2.2.1 Acquisition of stop-signal EEG and performance data (hypotheses 1, 2, 3)

See Kenemans et al. (2022) for details of the experimental procedures regarding the acquisition of the data relevant for hypotheses 1 and 2. The stop task took about 1 h and 30 minutes and subjects were offered a break halfway. EEG was recorded during the task.

#### 2.2.2 Acquisition of resting state EEG and reward-related performance data (hypothesis 4, 5)

For the Schutte et al. (2019) sample (Schutte et al., 2019) the 4-min resting state EEG was either recorded during the same session as the cued Go/No Go (CGN) task (always before the task or any other task performance) or it was recorded in a prior test session before any other task performance. Participants of the Schutte et al. (2020) sample (Schutte et al., 2020) attended three sessions during which they either received a dopaminergic drug, a noradrenergic drug, or a placebo. For the current study we only used data from the placebo session and only included subjects who received the placebo during the first session. These participants were first subjected to a simple 10-min motor task (not reported here), followed by a short version of the task (not reported here) and placebo administration. Resting state EEG data were recorded after a 2 h 40 min break. Subsequently, the CGN task was administered.

Thus, for all participants included in the analysis of hypothesis 3 resting state EEG data were collected before the start of the CGN task. All participants sat in a chair 1 meter in front of a computer screen in a separate dimly lit room next to the control room.

#### 2.2.3 Acquisition of resting state EEG and auditory stop-signal performance data (hypothesis 6)

Details of the experimental procedure can be found in Massar (2012). Participants were seated in a dimly lit room at a distance from the computer screen of approximately 60 cm. EEG signals were recorded during the task and participants filled in questionnaires at the end of the experimental session.

### 2.3 Behavioral tasks

#### 2.3.1 Stop signal task

Stopping performance was assessed by means of a stop signal task. The stop task used for the analysis of hypothesis 1, 2, and 3 (anticipated relationship between alpha, impulse control, and salience manipulation) is summarized below. Details can be found in (Kenemans et al., 2022). Subjects discriminated between two letter stimuli (go stimuli; “X” and “O”) by pressing a pre-specified button. Go stimuli were presented slightly above a fixation cross for 150 ms. Inter-trial intervals were set between 1500 and 1800 milliseconds.

Twenty-five percent of go stimuli were followed by a stop signal. The stop signal indicated that subjects had to try to withhold their already ongoing response to the go stimulus. The salience of the stop signal was manipulated across three different conditions: an auditory, and a high and a low-salient visual stop signal condition. In the auditory condition the stop signal was a 1000-Hz tone played binaurally at approximately 70 dB for 150 ms. The highly salient stop signal was a red background of the computer screen with a duration of 150 ms. The low-salient stop signal was a dollar sign with a visual angle comparable to the go stimuli.

The task consisted of 1 practice block (go stimulus only) with 126 trials which was followed by the three salience conditions. The order of the conditions was counterbalanced across subjects. Each salience condition started with a practice block followed by three main blocks, each consisting of 128 trials. After three main blocks the stimulus-hand mapping was switched and another practice block and three main blocks followed. The practice blocks also served to determine the duration of the optimal (i.e., yielding approximately 50 % successful stops) average go-stop interval of the first main block of a particular condition. The initial go-stop interval of each practise block was set to be the average go-stop interval of the previous main block, after applying an adaptive tracking algorithm (De Jong, Coles, & Logan, 1995). The go-stop interval of the first trial of the very first practice block was set to 250 ms. During all practice blocks go-stop intervals were increased by 50 ms after a successful stop, and shortened by 50 ms after a failed stop trial. The average go-stop intervals for main blocks 2 and 3 were based on the stopping performance of the previous block and determined by means of an adaptive tracking algorithm (De Jong et al., 1995). For all main blocks, go-stop intervals were jittered from -99 ms to 99 ms relative to the average go-stop interval.

Participants included in the analysis of the relation between resting state theta/ beta power ratio and stopping competence were subjected to only the auditory-stop version of a similar stop task. Details can be found in Massar (2012). Go stimuli in this task were blue circles and squares. Fourty percent of stop stimuli were followed by a stop signal, which consisted of a 1000-Hz tone played at 83 dB for 400 ms. Average go-stop intervals for each block were based on a staircase procedure (Massar, 2012). Go-stop intervals were jittered from -125 ms to 125 ms relative to the average go-stop interval. Subjects performed 8 task blocks consisting of 126 trials each. After 4 task blocks stimulus-response hand mapping was switched.

#### 2.3.2 CGN task

Details of the CGN task are described elsewhere (Schutte et al., 2019). In short, letters were presented on the screen and participants were instructed to press the right button when a cue letter (always letter A) was followed by target letter X and to press the left button when cue letter A was followed by target letter Y, as quickly and accurately as possible. The response hand-target mapping was counterbalanced across participants. Each letter stimulus was presented for 150 ms and was followed by an inter-stimulus interval with a random duration between 1400-1600 ms. In addition to A, X, and Y, the sequence was made up of 8 other letters, and the overall probability of an A was 20%.

The CGN task consisted of a practice block with 100 letter presentations and four main blocks of 400 letter presentations. The blocks differed with respect to whether correct and fast responding was rewarded or not and with respect to the probability of target appearance after the cue. The four blocks covered the following conditions: 0 Euros/ 50% probability given A, 0 Euros/ 98% probability given A, a maximum of 5 Euros/ 50% probability given A, and a maximum of 5 Euros/ 98% probability given A. The order of these conditions was counterbalanced across participants and participants were fully informed about the reward and probability condition of a block before the start of the block.

### 2.4 Resting state EEG

All resting state EEG data were recorded using the Active-Two system (Biosemi, Amsterdam, The Netherlands). Sixty-four electrodes were placed according to the international 10:10 system. The online sampling rate was 2048 Hz and data were filtered online with a default 400 Hz low-pass filter. Two-minutes of resting state EEG data were collected during an eyes-open condition and two-minutes of EEG data were collected during an eyes-closed condition.

### 2.5 Data reduction and statistical analysis

#### 2.5.1 Stopping performance data

For hypothesis 1-3, Stop signal reaction times (SSRTs) were estimated by means of the integration method as described in Verbruggen et al. (2019). Successful stops were defined as stop trials not followed by a response and the proportion of successful stops was computed. Reaction times (RTs) of all go trials (including omission trials and trials with pre-mature responses; RTs of trials with omissions were set to 1500 ms) were rank-ordered from shortest to longest for each block. The timing of the end of the stop process is estimated to correspond to reaction time *n* in the rank-ordered RT distribution. *N* is computed by multiplying the number of go trial RTs in that distribution by the probability of a failed stop during a given block (i.e., 1-proportion successful stops). SSRTs were computed for each block and condition separately by subtraction of the average go-stop interval of a given block from the *N*th RT of that block. SSRTs were subsequently averaged across the 6 blocks for each of the three salience conditions.

High-minus-low salience difference scores were created by subtraction of SSRTs for the low-salient stop signal from those for the high-salient and auditory conditions, respectively (i.e., SSRT high-salient minus SSRT low-salient and SSRT auditory minus SSRT low-salient).

For hypothesis 6 we used the same procedure to estimate SSRTs as used in the Lansbergen et al. study (Lansbergen et al., 2007), previously described in (De Jong, Coles, Logan, & Gratton, 1990). This method differs from the integration method in that the RT distribution only includes valid responses to go stimuli (trials with RTs between 150-1500 ms after stimulus onset and omission trials were excluded). Furthermore, the proportion of successful stops was corrected for omissions using the procedure described in (Tannock, Schachar, Carr, Chajczyk, & Logan, 1989).

#### 2.5.2 CGN task performance

Mean reaction times (RTs) for single responses to the target in the CGN task within a time window of 100-1500 ms after target onset were computed for each of the four reward/ probability conditions and for each subject. Furthermore the percentage correct responses to the target was computed for each of the four conditions and for each subject. RT and % correct data were subsequently averaged across the 50 and 98 % target probability condition yielding one value for the reward condition and one value for the no reward condition for each of the variables and for each subject. Reward-related performance scores were computed by subtracting the RT of the reward from the no reward condition and subtracting the % correct of the no reward from the reward condition.

#### 2.5.3 Stop task-related EEG analysis

Stop task-related EEG data were analyzed offline using Brainvision Analyzer version 2.1.2.327 (Brain Products, GmbH). Data were re-referenced to the signal of the right mastoid and the sampling rate was set to 250 Hz. A 0.5 Hz high-pass and notch filter were applied subsequently. Data were segmented from -950 until +50 ms surrounding go stimuli that were followed by a stop signal. Data were segmented separately for go stimuli followed by correct and failed stops. Before eye blink correction we removed segments with extreme artifacts (maximal allowed voltage step: 50 μV, amplitude between -400 and +400 μV, and lowest allowed activity: 0.5 μV). This was done in order to improve the Gratton and Coles ocular correction method (Gratton, Coles, & Donchin, 1983). Remaining artifacts after the ocular correction were removed for the target electrodes POz and Pz at which alpha power was maximal (maximal allowed difference of values: 200 μV). For each subject the numbers of remaining epochs for the successful and failed stop trials were equalized by a random removal procedure.

Spectral power in the alpha band was estimated by means of a Fast Fourier transformation with a 10% Hanning window. The apriori selection of the frequency band (9-10.5 Hz) and target electrodes (POz, Pz) was based on a collapsed localizer approach. Alpha power was averaged across the two target electrodes and averaged across all stop trials of the low-salient visual condition. Figure 1 displays the distribution of alpha power across the scalp.

**Fig 1.**
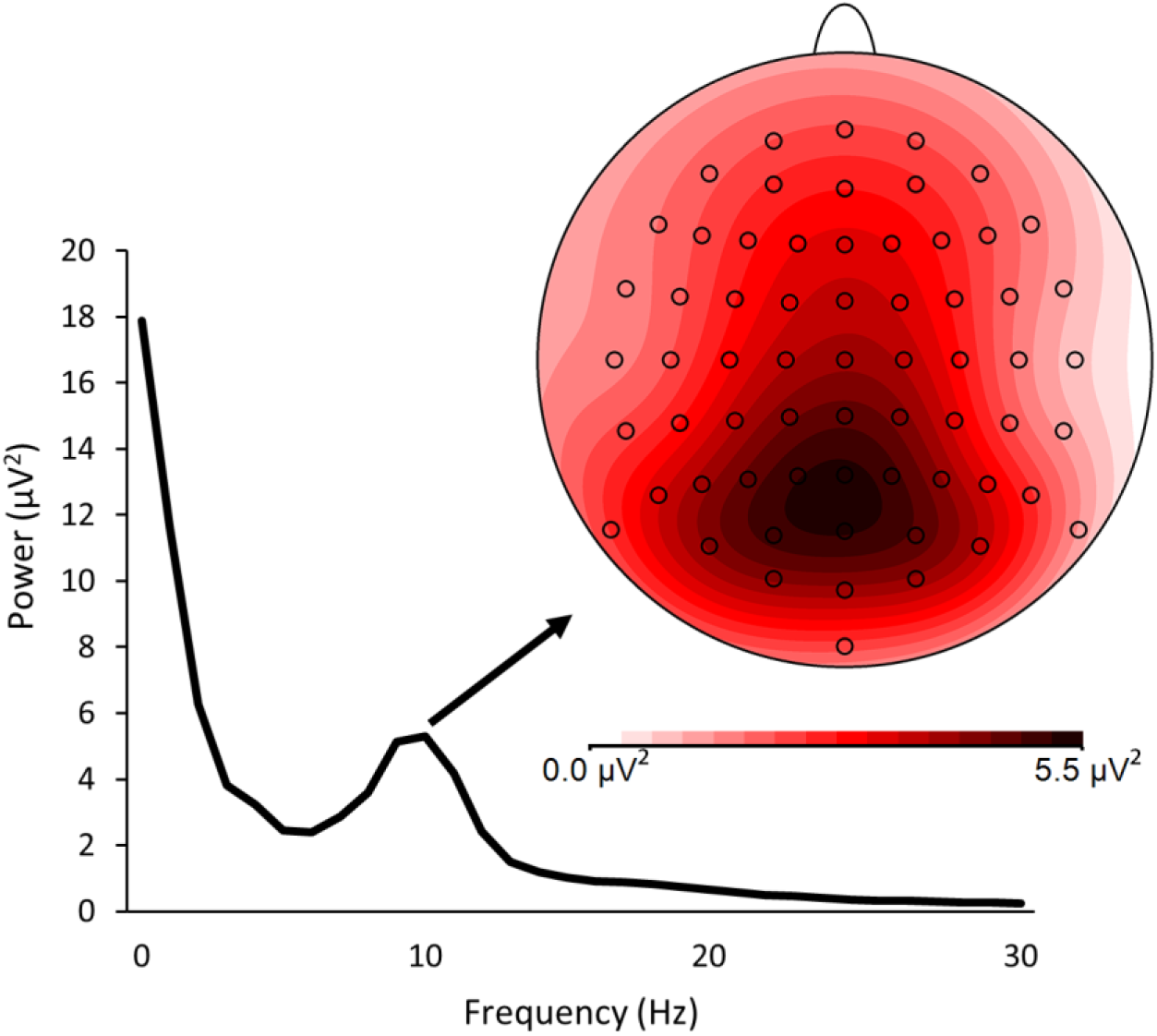
Power spectrum and distribution of pre-stop stimulus alpha power across the scalp averaged across subjects. The figure displays the power spectrum (lower left) averaged across electrode Pz and POz for frequencies between 0 and 30 Hz. The figure corresponds to the stop task-related data set. Pre-stop stimulus alpha power had a peak frequency of 9.8 Hz and was maximal over the parieto-occipital cortex (scalp plot; upper right).

#### 2.5.4 Resting-state EEG data analysis

All resting state EEG data were analyzed offline using Brainvision Analyzer version 2.1.2.327 (Brain Products, GmbH). The analysis steps for examining the anticipated relationship between theta/ beta power ratio and stopping competence, and those for association between theta/ beta power ratio, and reward-related performance were similar, unless stated otherwise. Sampling rate was set to 256 Hz and the averaged signal of the mastoids was used as reference. Data were filtered using a 1-40 Hz bandpass filter at 12 dB/oct. Data were divided into 2-sec segments for the eyes open and eyes closed condition separately and corrected for ocular artifacts using the Gratton & Coles algorithm (Gratton et al., 1983). A baseline correction was applied subsequently to suppress potential DC drifts. Channels of interest (Fz, FCz) were individually inspected for artifacts and segments with data points exceeding 50 or -50 microvolts were automatically removed.

With respect to the theta/ beta-stop performance resting state data, spectral power in the theta (4-7 Hz) and beta (13-30 Hz) band were estimated by means of a Fast Fourier transformation with a 10% Hanning window. Spectral power values were exported for electrode FCz at which the theta/ beta power ratio across subjects was maximal. Spectral power values were averaged across segments for the eyes open and eyes closed condition separately and theta/ beta power ratios were computed. Data of one subject were removed from the analysis because of a procedural error during the experiment. Figure 2 displays the power spectrum and scalp distribution of theta/ beta power ratio.

**Fig 2.**
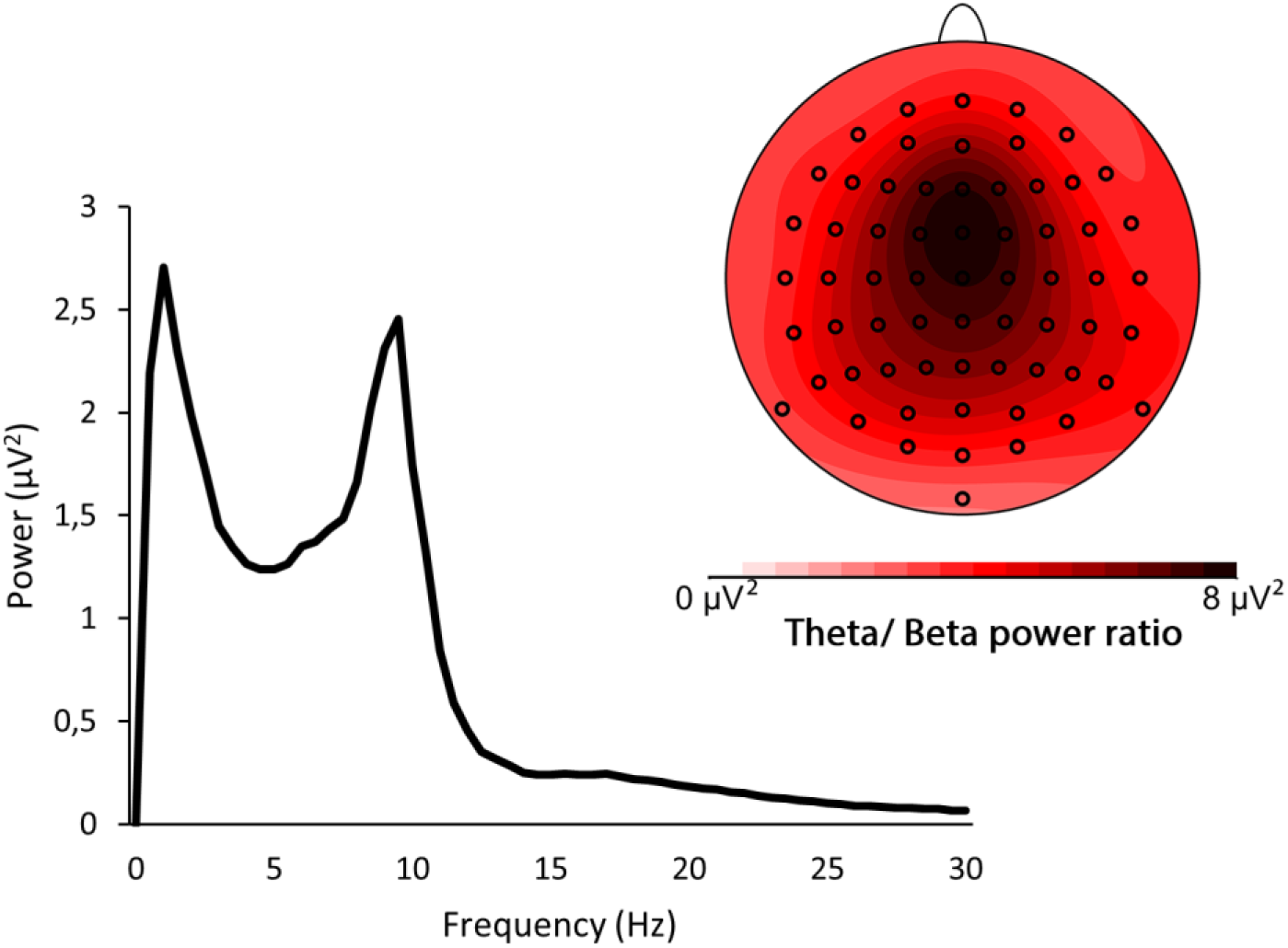
Power spectrum and distribution of resting-state theta/ beta power ratio across the scalp averaged across subjects (theta/ beta-stop performance). The figure displays the power spectrum (lower left) for electrode FCz for frequencies between 0 and 30 Hz and corresponds to the theta/ beta-stop performance resting-state data set (the anticipated relation between resting state theta/ beta power ratio and stop performance). Data were collapsed across the eyes open and eyes closed condition. Theta/ beta power ratio was maximal over the frontal-central cortex (scalp plot; upper right). In addition to the artifact removal procedure described in section 2.5.4, all channels not of primary interest were (automatically) inspected for artifacts and segments with data points exceeding -100 μV or 100 μV were removed for these channels. This was done to reduce noise for visualization purposes. For channels F3, F4, F7, and F8 the artifact criterium was set to be the same as for the channels of primary interest (Fz, FCz).

With respect to the theta/ beta-reward resting state data, spectral power values in the theta (4-7 Hz) and beta (13-30 Hz) band were exported for electrode FCz at which the theta/ beta power ratio across subjects was maximal. Figure 3 displays the power spectrum and scalp distribution of the theta/ beta power ratio for the theta/ beta-reward data set. Spectral power values were averaged across segments for the eyes open and eyes closed condition separately, and theta/ beta power ratios were computed. One subject was removed from the analysis due to a technical failure.

**Fig 3.**
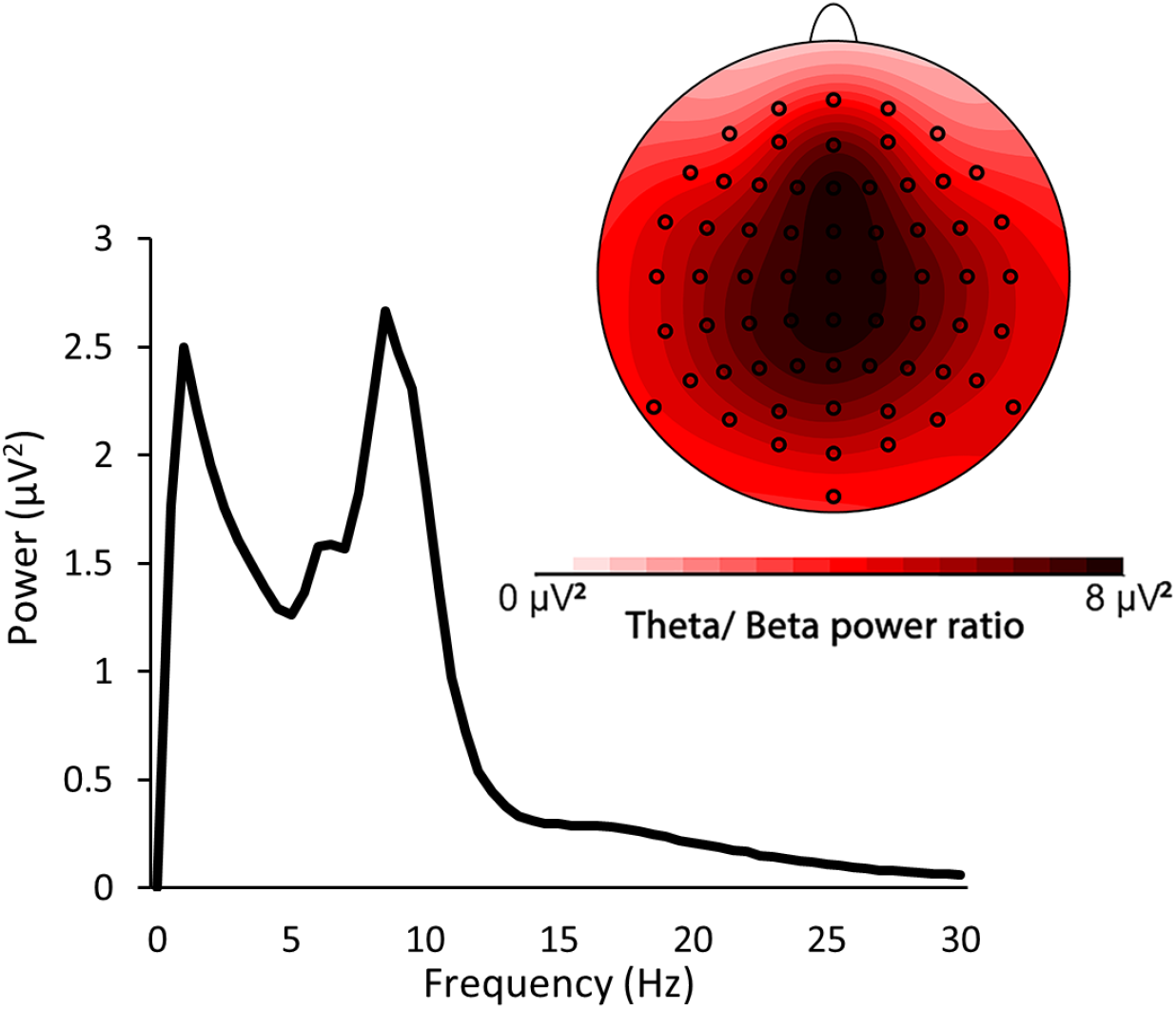
Power spectrum and distribution of resting-state theta/ beta power ratio across the scalp averaged across subjects (theta/ beta-reward-related performance). The figure displays the power spectrum (lower left) for electrode FCz for frequencies between 0 and 30 Hz and corresponds to the theta/ beta-reward-related performance resting-state data set (the anticipated relation between resting state theta/ beta ratio and reward-related performance). Data were collapsed across the eyes open and eyes closed condition. Theta/ beta power ratio was maximal over the frontal-central cortex. In addition to the artifact removal procedure described in section 2.5.4, all channels not of primary interest were (automatically) inspected for artifacts and segments with data points exceeding -100 μV or +100 μV were removed for these channels. This was done to reduce noise for visualization purposes. For channels F3, F4, F7, and F8 the artifact criterium was set to be the same as for the channels of primary interest (Fz, FCz). Data of four subjects were not included in the scalp plot, because of one or more noisy channels with less than half of the segments left.

#### 2.5.5 Statistical analysis

Absolute theta and beta band power during the resting state eyes-open condition was highly correlated with theta and beta power during the eyes-closed condition, all correlation coefficients ≥ .76. All resting state EEG data were therefore collapsed across the eyes-open and eyes-closed conditions.

For all analyses, raw variables or transformed variables (contrasts) were tested for normality (Shapiro-Wilk). Deviation of normality prompted non-parametric testing as specified below.

A non-parametric Spearman’s correlation test was performed to test the relationship between pre-stimulus alpha power on the one hand, and stopping performance on the other hand.

A mixed multivariate analysis of variance (MANOVA) was performed with stop-stimulus salience (low-salient versus high-salient versus auditory) as within-factor and median-split based alpha power as between-subjects factor to test effects of salience on stopping performance, and whether these differed between low- and high-susceptibility (high versus low alpha power) individuals. In case of multivariate salience effects, pairwise comparisons between salience conditions were made using either t tests or Wilcoxon-z tests.

As reward-non-reward contrasts (for RT and error rate) were non-normally distributed in both theta/beta groups, effects of reward, and how these differ between theta/beta groups, were tested using Wilcoxon rank tests and Mann-Whitney U tests, respectively.

A non-parametric Mann-Whitney U test was applied to the difference in stopping competence between the high and low resting-state theta/ beta groups.

Alpha was set to .05. For all post-hoc comparisons testing the effect of salience or reward for groups separately, alpha was set to .025 in order to correct for two within-group comparisons (Bonferroni correction).

## 3. Results

### 3.1 Relationship between EEG pre-stop stimulus alpha and stopping competence (hypothesis 1)

Low pre-stop stimulus alpha power (9-10.5 Hz) was not significantly related to better stopping, rho(28) = .203, *p* = .283.

### 3.2 The interaction between EEG pre-stop stimulus alpha and the effect of salience on stopping competence (hypothesis 2-3)

On average stop-signal salience had a significant impact on stop signal reaction times, F(2,27) = 26.36, *p* < .001. Post-hoc pairwise comparisons showed that SSRTs were significantly longer when the stop-signal was low-compared to high-salient (i.e., a high-salient visual or auditory signal), t(29) = 7.45, *p* < .001 for low-salient compared to auditory and Z = -3.86, *p* < .001 for low-salient compared to salient. SSRTs were also significantly longer for the high-salient visual condition compared to the auditory stop-signal condition, t(29) = 3.69, *p* < .001. This result indicates that stopping performance improved (i.e., shorter SSRTs) with increasing stop-signal salience (i.e., performance low-salient visual < high-salient visual < auditory). The effect of stop-signal salience on SSRTs was, however, not significantly different between participants with high and low median-split based alpha power, F(2,27) = .22, *p* = .806. Figure 4 presents the SSRTs for each stop-signal salience condition and for each alpha power group separately.

**Fig 4.**
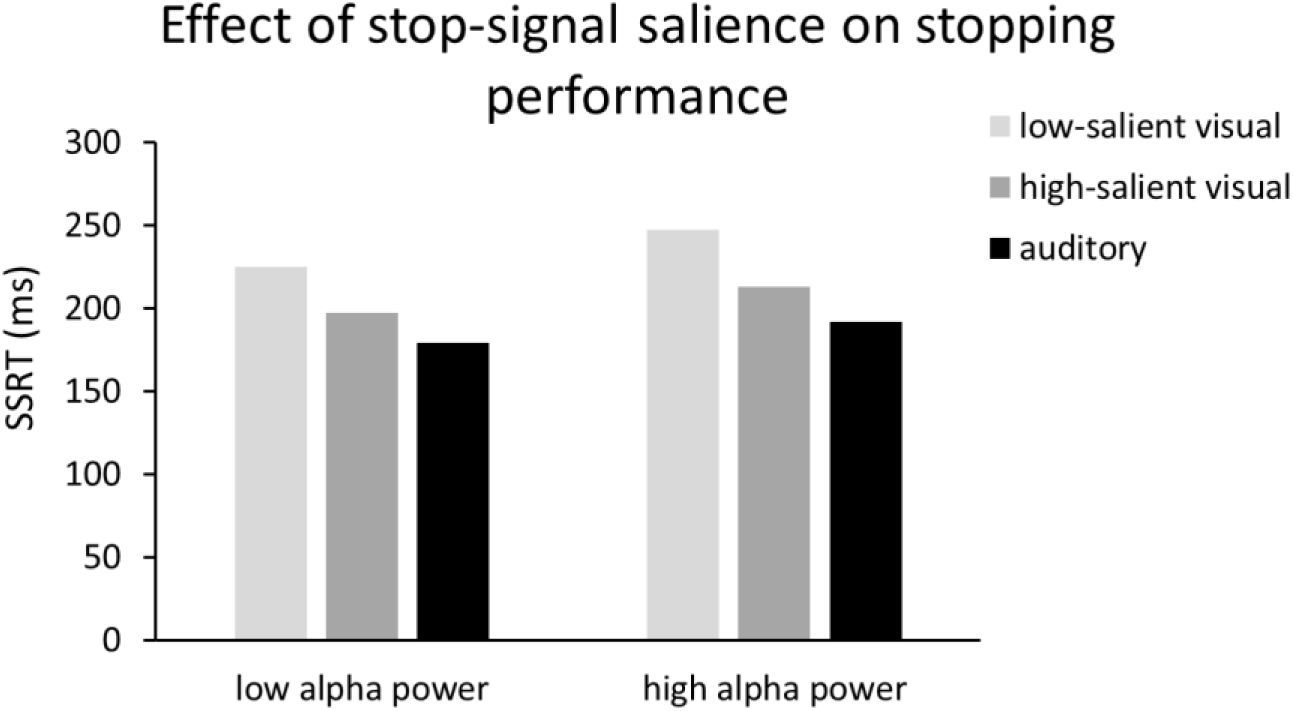
The effect of stop-signal salience on stopping performance in individuals with low and high pre-stop stimulus alpha power. On average stop signal reaction times were significantly higher in the low- (light grey bars) compared to the high-salient visual (dark grey bars) and auditory (black bars) stop-signal conditions. This effect of salience on stopping performance was not significantly different between participants with low and high pre-stimulus alpha power (9-10.5 Hz).

### 3.3 The interaction between EEG theta / beta power and reward-related performance (hypothesis 4-5)

On average participants responded faster and showed more correct responses during reward blocks (mdn RT = 463 ms) compared to no-reward blocks (mdn RT = 483 ms), as shown by significant Wilcoxon signed-rank tests, *Z* = -4.25, *p* < .001, *Z* = -3.07, *p* = .002, respectively. The effect of reward on correct responding was significantly different between participants with a high and low midfrontal theta/ beta power ratio, *Z* (Mann Whitney) = -2.42, *p* = .015. Non-parametric (Wilcoxon) follow-up tests for the high and low theta/ beta group separately showed that the effect of reward on accuracy (i.e., % correct reward minus no reward) was only significant in the low theta/ beta group, *Z* = -3.46, *p* < .001, and not in the high theta/ beta group, *Z* = -0.7, *p* = .486. This result indicates that individuals with a relatively low resting state midfrontal theta/ beta power ratio showed a larger increase in correct responses when a reward could be obtained as compared to individuals with a high resting state frontal theta/ beta power ratio (Figure 5).

**Fig 5.**
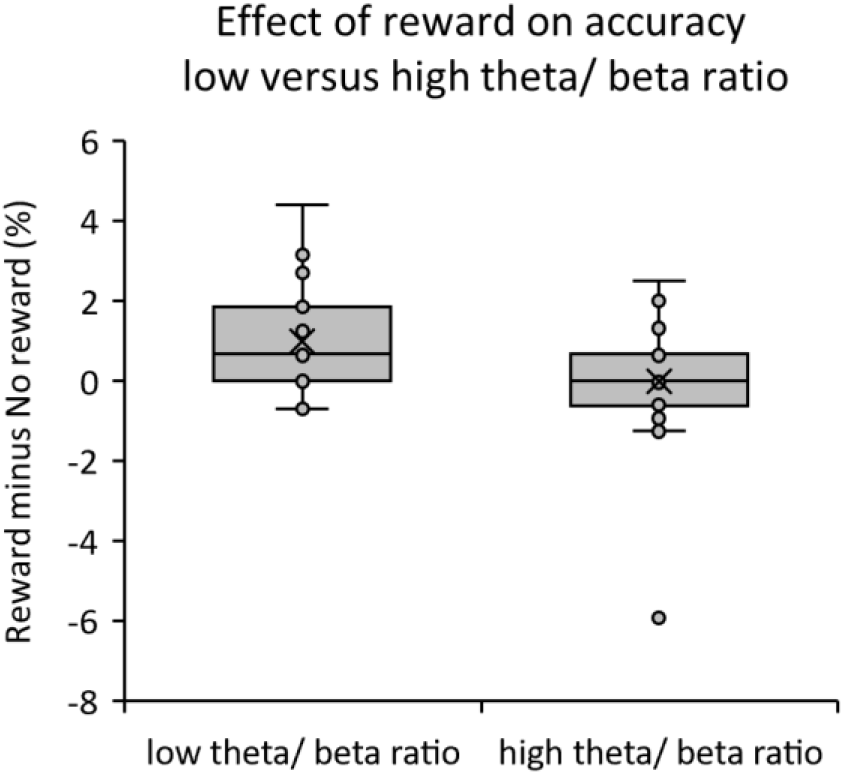
The effect of reward on accuracy in participants with a low and high theta/ beta power ratio. On average participants with a low median-split based theta/ beta power ratio showed more correct responses during reward compared to no reward blocks. No such effect was observed for participants with a high median-split based theta/ beta power ratio.

There was no difference between theta/ beta power ratio groups with respect to the reward-related decrease in reaction times, *Z* = -1.03, *p* = .304.

### 3.4 The effect of resting state EEG theta/ beta power ratio on stopping competence (hypothesis 6; replication experiment)

Stopping competence was not significantly different between subjects with high (mdn = 188 ms) and low (mdn = 193 ms) median-split based resting state theta/ beta power ratios, *U* (N_low_= 26, N_high_= 25) = 320, *Z* = -.094, p = .925.

## 4. Discussion

The current study tested whether impulse control can be enhanced by interventions regarding the salience of external cues that signal the need for impulse control and regarding reward. In addition, we investigated how individual characteristics can contribute to differences in responsivity to such interventions. This was adressed by relating putative markers of environmental susceptibility (EEG alpha power) and reward sensitivity (theta/ beta power ratio) to impulse control and reward-induced performance enhancement, respectively. These electrophysiological markers, in turn, were used as a means to examine individual differences in responsivity to the reward and salience interventions.

Our results show that impulse control can be enhanced simply by increasing the salience of external cues. We did, however, not find evidence for a relationship between EEG alpha power and impulse control, and no differences in responsivity to the salience intervention between participants with high and low alpha power. We found that introducing a reward for correct performance induces overall better task performance. Moreover, we found evidence for individual differences in responsivity to the reward intervention. In participants with a relatively low theta/ beta power ratio, reward increased correct action selection.

Prior studies (Dockree et al., 2017; O’Connell et al., 2009) have shown that EEG alpha power preceding a visual target is associated with detection errors, suggesting that EEG alpha power can be seen as an electrophysiological correlate for the susceptibility to signals in the environment. Therefore we hypothesized that participants with relatively low levels of alpha power preceding stop signals would be better able to attend to or to process stop signals. Hence, it was expected that low levels of alpha power preceding stop signals would be associated with enhanced stopping performance, the objective index of impulse control (Verbruggen et al., 2019). We anticipated that increasing the salience of stop signals (i.e., making them more distinguishable) would lead to enhanced stopping performance and that this intervention would have a greater effect in participants with high alpha power. The salience manipulation did exert the expected effect on overall stopping performance. In line with previous studies stopping was faster when the stop signal was high compared to low in salience. (Blizzard et al., 2017; Montanari et al., 2017; Van Der Schoot et al., 2005). Contrary to our expectation, we did not observe a relationship between EEG alpha power preceding stop signals and stopping performance, and no difference in the effect of the salience manipulation between participants with high and low alpha power. These findings indicate that enhancement of the salience of external cues is an effective intervention for the overall improvement of impulse control. This result also suggests that average task-related alpha power may not reflect trait-like susceptibility to signals in the environment. Results of studies showing that alpha power can distinguish between misses and hits (Dockree et al., 2017; O’Connell et al., 2009) may rather indicate that alpha power fluctuations across trials result in transient shifts in susceptibility.

In a prior study it was found that alpha power attenuation from an eyes-closed to eyes-open resting-state condition (alpha reactivity) was larger for individuals diagnosed with ADHD compared to control subjects. No such effect was observed for absolute alpha power during either the eyes-open or eyes-closed condition (van Dongen-Boomsma et al., 2010). Based on these findings it is possible that, unlike task-related alpha power, alpha reactivity during resting state conditions relates to stopping performance and the effect of the salience manipulation on stopping. This may be an interesting avenue for future research.

Several studies have shown that reward affects performance in go/ no go tasks, e.g., (Demurie, Roeyers, Wiersema, & Sonuga-Barke, 2016; Schutte et al., 2019). This performance improvement by reward is likely due to interactions between reward-related brain areas and networks related to perception and executive control (Pessoa & Engelmann, 2010; Schutte et al., 2019). Several studies have also shown that reward improves stopping performance (Boehler et al., 2014; Demurie et al., 2016; Doekemeijer, Verbruggen, & Boehler, 2021; Kohls, Herpertz-Dahlmann, & Konrad, 2009), indicating that rewarding successful impulse control may be a promising intervention strategy to improve impulse control.

Prior studies have shown that frontal theta/ beta power ratio is positively related to reward-learning, risky decision-making, and reward-seeking during gambling tasks (Kelley, Hortensius, Schutter, & Harmon-Jones, 2017; Massar et al., 2014; Schutter & Van Honk, 2005). In an earlier study it was found that stopping performance was better in individuals with high theta/ beta power ratio’s, which in turn was interpreted as reflecting stronger task-induced motivation in the high theta/ beta group. Hence, individuals with low theta/ beta power ratio were expected to benefit more an extrinsic monetary reward.

The results show that the reward intervention did exert the expected improving effect on overall performance during the cued go/ no go task. More specifically, in line with our hypothesis, the response to the reward intervention was enhanced in participants with a relatively low theta/ beta power ratio. These individuals showed more correct responses when a reward compared to no reward could be obtained compared to individuals with a relatively high theta/ beta power ratio. This finding is consistent with our hypotheses that theta/ beta power ratio is associated with task-induced motivation and that an extrinsic reward intervention exerts a greater effect on performance in individuals with weak task-induced motivation. Individuals with a low theta/ beta power ratio may have had lower levels of intrinsic motivation to perform well on the task in general (Lansbergen et al., 2007). Consequently, the addition of performance-contingent reward may have led to improved performance specifically in the low theta/ beta group, because there was more room for improvement to occur in this group compared to the high theta/ beta power ratio group. The high theta/ beta group may have had higher levels of motivation to perform well, even without additional reward (Lansbergen et al., 2007). An alternative interpretation is that individuals with a relatively low theta/ beta power ratio showed more correct responses during reward blocks, because they were more sensitive to punishment (which results from an incorrect response during a reward block). A prior study, however, has shown that the positive relationship between theta/ beta power ratio and risky decision-making during the Iowa gambling task is driven by these individuals displaying increased approach rather than reduced avoidance behavior (Massar et al., 2014). This finding renders the alternative interpretation less likely.

It has been argued that the theta/ beta power ratio reflects cortical-subcortical interaction and as such the theta/ beta power ratio may involve the extent to which reward (subcortex) modulates cognitive control and attention (cortex) (Schutter & Knyazev, 2012) or vice versa (Angelidis, Hagenaars, van Son, van der Does, & Putman, 2018). Results of prior studies have shown that a low resting-state theta/ beta power ratio is associated with high resilience against disruption of attentional control by emotional stimuli (Angelidis et al., 2018). These latter results suggest that theta/ beta power ratio can also be seen as a neural marker of attentional control over emotional stimuli, rather than as a neural marker of reward sensitivity. Interestingly, the results of the current study show that the high and low theta/ beta power ratio groups only differed with respect to the effect of the reward intervention on accuracy, not with respect to the effect of the reward intervention on reaction speed. It may be speculated that individuals with a relatively low theta/ beta power ratio were better able to adjust the level of cognitive control in order to improve the accuracy of their responses during the reward blocks; this idea is consistent with thet results of a recent tACS study (Wischnewski, Joergensen, Compen, & Schutter, 2020).

Finally, it should be noted that in a previous study (Lansbergen et al., 2007) it was reported that individuals with a high theta/ beta power ratio during rest are better at stopping, this relationship was not replicated in the current study. One possible explanation for this discrepancy may be that in the earlier study only individuals scoring very high and very low on self-reported impulsivity were included. This is clearly deviant from the current study in which participants were not pre-selected. Furthermore, other findings have demonstrated a positive relationship between impulsivity and theta (Snyder & Hall, 2006) and a negative relationship between impulsivity (ADHD) and inhibitory control (stopping) (Lijffijt, Kenemans, Verbaten, & van Engeland, 2005). It is therefore possible that the difference in subject characteristics contributes to the discrepancy between our current finding and that of the earlier study.

In conclusion, results show that increasing the salience of external cues may be a feasible way to positively affect impulse control. The efficacy of reward interventions however partially depends on individual characteristics with respect to reward seeking tendencies. Reward interventions may be specifically effective in individuals with low levels of intrinsic motivation. From a practical point of view, the results imply that salience and reward interventions are possibly worthwhile as a measure to enhance adherence to rules such as the physical distancing rules during the Covid pandemic and that require inhibition of automatic behavioral tendencies and impulses. Examples of possible interventions are flags or salient auditory signals when a 1.5-meter boundary is transgressed or additional discounts in a supermarket after physical distancing.

## Acknowledgements

This study was funded by a Covid-19 research grant from the Utrecht University Faculty of Social and Behavioral Sciences awarded to LK and DS.

